# Variation of length and sequence of the nuclear ribosomal DNA internal transcribed spacer 1 supports “hermit-to-king” crab hypothesis

**DOI:** 10.1101/2022.07.24.501315

**Authors:** Seinen Chow, Katsuyuki Hamasaki, Kooichi Konishi, Takashi Yanagimoto, Ryota Wagatsuma, Haruko Takeyama

## Abstract

Lithodoid and paguroid crabs are morphologically assigned to the superfamilies Lithodoidea and Paguroidea, respectively. Molecular analyses, however, have revealed closer genetic proximity of the lithodoid crabs to the family Paguridae than to other families of Paguroidea, provoking a long debate. We investigated the length and nucleotide sequence variation of the nuclear ribosomal DNA internal transcribed spacer 1 (ITS1) in lithodoid and paguroid species. Uniquely short ITS1s (215–253 bp) were observed in seven lithodoid species belonging to the families Lithodidae and Hapalogastridae. In contrast, ITS1 length varied considerably in 13 paguroid species belonging to the families Coenobitidae, Diogenidae, and Paguridae. Short-to-long ITS1s (238–1090 bp) were observed in five species of the family Paguridae, and medium to long ITS1s (573–1166 bp) were observed in eight species of the families Coenobitidae and Diogenidae. Interestingly, ITS1s of considerably different sizes coexist in individual paguroid species. Nucleotide sequence analysis indicated that the short ITS1s observed in the family Paguridae were descendants of longer ITS1s and were homologous to the short ITS1 of lithodoid species. ITS1 sequences of the families Coenobitidae and Diogenidae shared no nucleotide elements similar to those of lithodoid and pagurid species. These molecular signals indicate that the short ITS1 in pagurid lineage was passed on to lithodoid lineage, strongly supporting the “hermit-to-king” crab hypothesis.

## 1 Introduction

The Anomura, one of the infraorders in Decapoda, Crustacea, is the morphologically and ecologically most diverse group. After numerous taxonomic revisions (for example, see McLaughlin et al. 2007, 2010), Anomura now comprises six extant superfamilies (Chirostyloidea, Galatheoidea, Hippoidea, Lithodoidea, Lomisoidea, and Paguroidea) (WoRMS 2022). Boas (1880) first pointed out that lithodid king crabs were modified hermit crabs of the genus *Pagurus* or free-living hermit crabs. This concept, called carcinization (Borradaile 1916) has generally been concordant with subsequent morphological studies on adults (McLaughlin 1983; but see Martin and Abele 1986) and larvae (MacDonald et al. 1957; Hart 1965; Lang and Young 1977; Campodonico and Guzmánm 1981; Haynes 1982, 1984; Konishi 1986). The first molecular approach by Cunningham et al. (1992) not only corroborated this carcinization concept but also suggested the inclusion of lithodid king crabs within the genus *Pagurus*—this is known as the “hermit-to-king” crab hypothesis. Richter and Scholtz (1994) and Scholtz (2014) presented the morphological characteristics of a hermit crab ancestry of lithodids, whereas McLaughlin and Lemaitre (1997), McLaughlin et al. (2004, 2007), and Lemaitre and McLaughlin (2009) developed theories against this hypothesis. Starting with Zaklin (2001), all subsequent studies using molecular genetic analyses have strongly supported the “hermit-to-king” crab hypothesis (Morrison et al. 2002; Ahyong and O’Meally 2004; Ahyong et al. 2009; Hall and Thatje 2009, 2018; Schnabel et al. 2011; Tsang et al. 2008; 2011; Bracken-Grissom et al. 2013; Noever and Glenner 2018; Tan et al. 2018, 2019). However, there is no consensus whether lithodids are nested within the genus *Pagurus*.

All molecular genetic studies mentioned above used mitochondrial DNA (mtDNA), nuclear protein-coding genes, and/or nuclear ribosomal RNA-encoding subunits (18S or 28S rDNA), while non-coding spacer regions flanked by the rRNA-encoding subunits were not utilized. Although the internal transcribed spacer 1 (ITS1) of nuclear ribosomal DNA (nrDNA) is usually divergent between species and homogenized within species, we observed large length and nucleotide sequence variations in ITS1 not only between but also within the species belonging to Lithodoidea and Paguroidea. Here, we analyzed the ITS1 of seven species of Lithodoidea and 13 species of Paguroidea.

## 2 Materials and Methods

The lithodoid and paguroid crabs collected in this study are listed in Table 1. In the superfamily Lithodoidea, three species (n = 5) from two genera (*Dermaturus* and *Hapalogaster*) of the family Hapalogastridae and four species (n = 4) from three genera (*Cryptolithodes, Lopholithodes*, and *Paralomis*) of the family Lithodidae were analyzed. In the superfamily Paguroidea, seven species (n = 9) from two genera (*Birgus* and *Coenobita*) of the family Coenobitidae, three species (n = 4) from three genera (*Aniculus, Areopaguristes*, and *Dardanus*) of the family Diogenidae, and five species (n = 7) from three genera (*Boninpagurus, Elassochirus*, and *Pagurus*) of the family Paguridae were analyzed. Four species (*Cryptolithodes expansus, Dermaturus mandtii, Hapalogaster grebnitzkii*, and *H. dentata*) were museum specimens stored in 70% ethanol, and others were frozen or fresh state specimens.

**Table 1.**
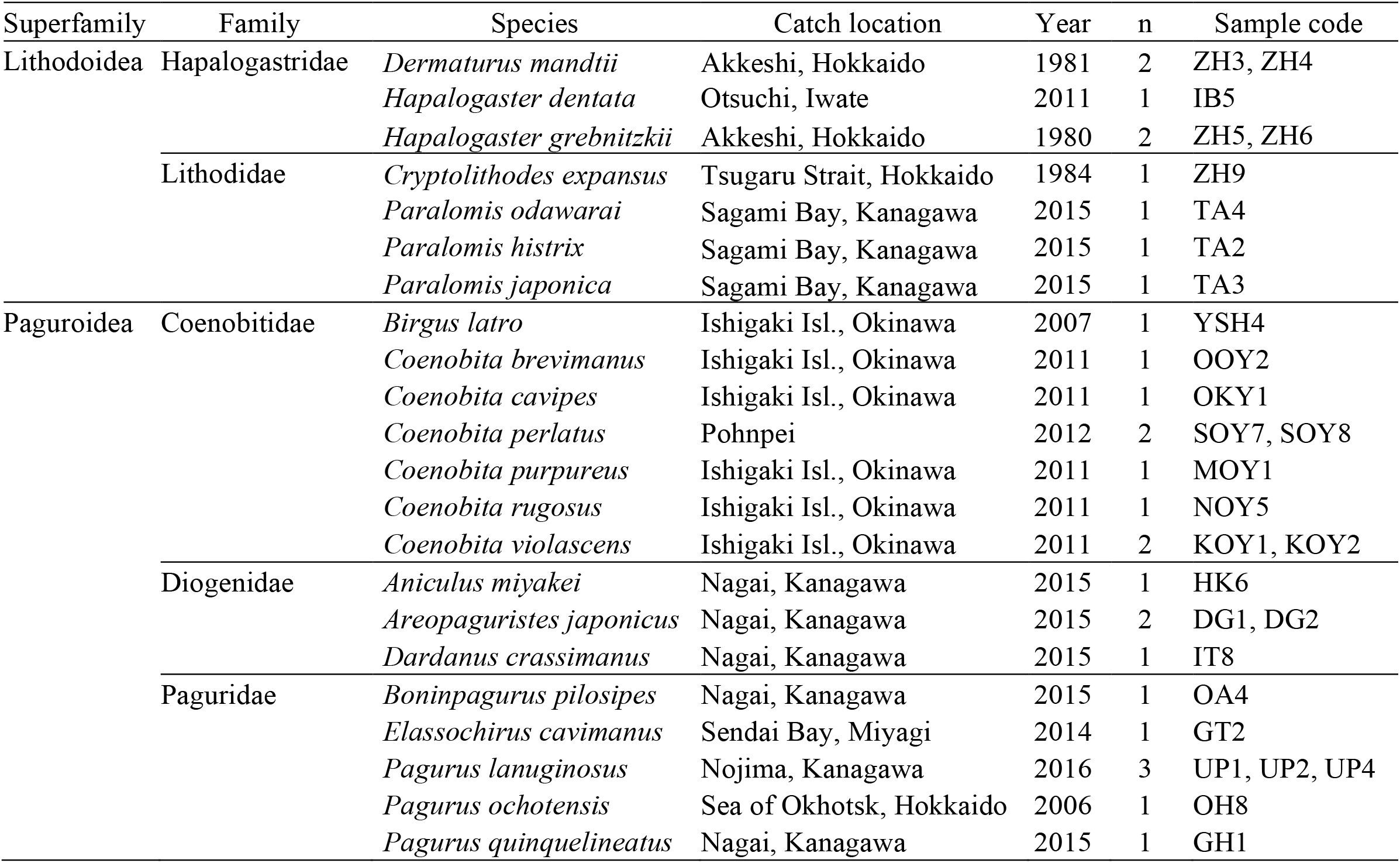
Sampling information of lithodoid and paguroid species used in the present study

Leg muscle tissue was used for DNA extraction using the QuickGene DNA tissue kit (DT-S, KURABO). The entire stretch of ribosomal DNA internal transcribed spacer 1 (ITS1) was amplified using a PCR primer pair (ITS1: TCCGTAGGTGAACCTGCGG; 5.8S: CGCTGCGTTCTTCATCG) (see Chow et al., 2009). PCR amplification was performed in a 12 μL reaction mixture containing 1 μL of template DNA (1-10 ng/μL), 1.2 μL of 10 × reaction buffer, 1 mM of each deoxynucleotide triphosphate, 0.4 μM of each primer, 0.5 U of EX Taq HS polymerase (Takara Bio, Inc.), and sterilized Milli-Q water. The reaction mixtures were preheated at 94 °C for 5 min, followed by 30 amplification cycles (denaturation at 94 °C for 0.5 min, annealing at 58 °C for 0.5 min, and extension at 72 °C for 1 min), with a final extension at 72 °C for 7 min. The PCR products were electrophoresed on a 1.5% agarose gel to confirm amplification. The PCR products were treated with ExoSAP-IT (GE Healthcare) to remove the PCR primers.

Direct nucleotide sequencing was performed using a BigDye Terminator Ver3.1 kit (Applied Biosystems) with forward and reverse PCR primers. Sequencing was conducted using an ABI3730XL automatic sequencer (Applied Biosystems). When sequence electropherograms generated by direct nucleotide sequencing were not readable, the PCR products were cloned using the DynaExpress TA PCR Cloning Kit (BioDynamics Laboratory Inc.). Colony-direct PCR was performed using M13 primers according to the reaction protocol described above, followed by agarose gel electrophoresis to confirm the size of the inserted fragments. Amplicons of different sizes observed within an individual or species were treated as described above to determine the nucleotide sequences.

All nucleotide sequences obtained were compared against the GENBANK database using the BLASTN program (Altschul et al. 1990) to identify similar sequences. Nucleotide sequences were aligned using CLUSTAL W (as implemented in GENETYX, GENETYX Inc., Tokyo), followed by manual editing. Selection of the best-fit model for nucleotide substitution and construction of phylogenetic trees were performed using MEGA 6 (Tamura et al. 2013).

## 3 Results

### 3.1 Overview of ITS1 sequences

Single short fragments (ca. 300 bp) were amplified from nine individuals of all seven lithodoid species examined, and direct nucleotide sequencing of the amplicons was unproblematic. In contrast, as shown in Fig. 1, amplification of multiple fragments was commonly observed in many paguroid species examined. Since we often failed to obtain good electropherograms using direct nucleotide sequencing, the PCR amplicons of these paguroid individuals were cloned and subjected to nucleotide sequencing. Clones with inserts of different sizes within an individual were selected and the nucleotide sequences of 35 clones from 13 paguroid species were successfully determined. We failed to obtain transformed clones from two species (*Coenobita perlatus* and *C. violascens*). All sequences determined (the International Nucleotide Sequence Database Collection: accession No. LC706586‒LC706620) were consisted of the 3′ end of 18S rDNA, 5′ end of 5.8S rDNA, and the complete ITS1 in between.

**Fig. 1.**
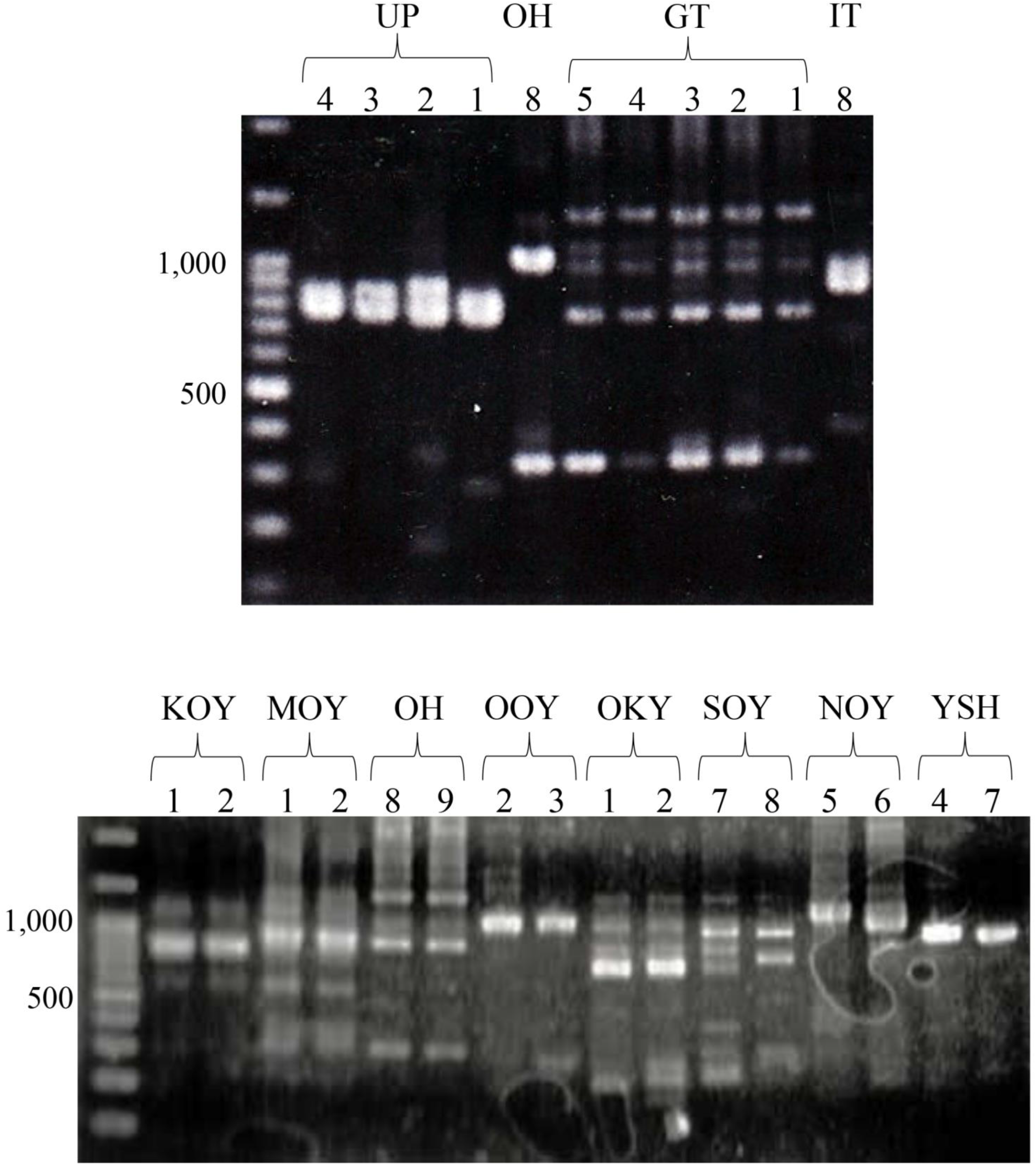
Agarose gel electrophoresis images of ITS1 amplicons of 10 paguroid species. Left ends are molecular markers. Coenobitidae (KOY: *Coenobita violascens*, MOY: *C. purpureus*, NOY: *C. rugosu* OKY: *C. cavipes*, OOY: *C. brevimanus*, SOY: *C. perlatus*, YSH: *Birgus latro*), and Paguridae (UP: *Pagurus lanuginosus*, OH: *P. ochotensis*, GT: *Elassochirus cavimanus*). See Table 1 for specimen number of each species

The length and GC content of ITS1, and the results of the BLAST search are presented in Table 2. According to length, ITS1 sequences were categorized into short (< 400 bp), medium (400 to 799 bp), and long (≥ 800 bp), and designated as S-, M-, and L-types, respectively. ITS1 sequences of all nine individuals of seven lithodoid species were S-type, ranging from 217 to 253 bp, and a BLAST search indicated these sequences to be homologous to ITS1s of lithodoid species in the database. Moreover, S-type ITS1 may be unique to the superfamily Lithodoidea, since all 53 ITS1 sequences of 19 king crab species of the family Lithodidae reported to date (HM020983-HM021023, AB194389-AB194394, AB211306, AB236928, AB426492-AB426495) (also see Chow et al. 2009; Hall and Thatje 2018) were also short (215–219 bp). Of the 26 nucleotide sequences obtained from 13 paguroid species examined in the present study, four ITS1s (LC706617‒ LC706620, 155–262 bp) were determined to belong to fungi or jellyfish (*Anemonia erythraea*) and were not included in the subsequent analysis as they were considered cross-contamination.

**Table 2.**
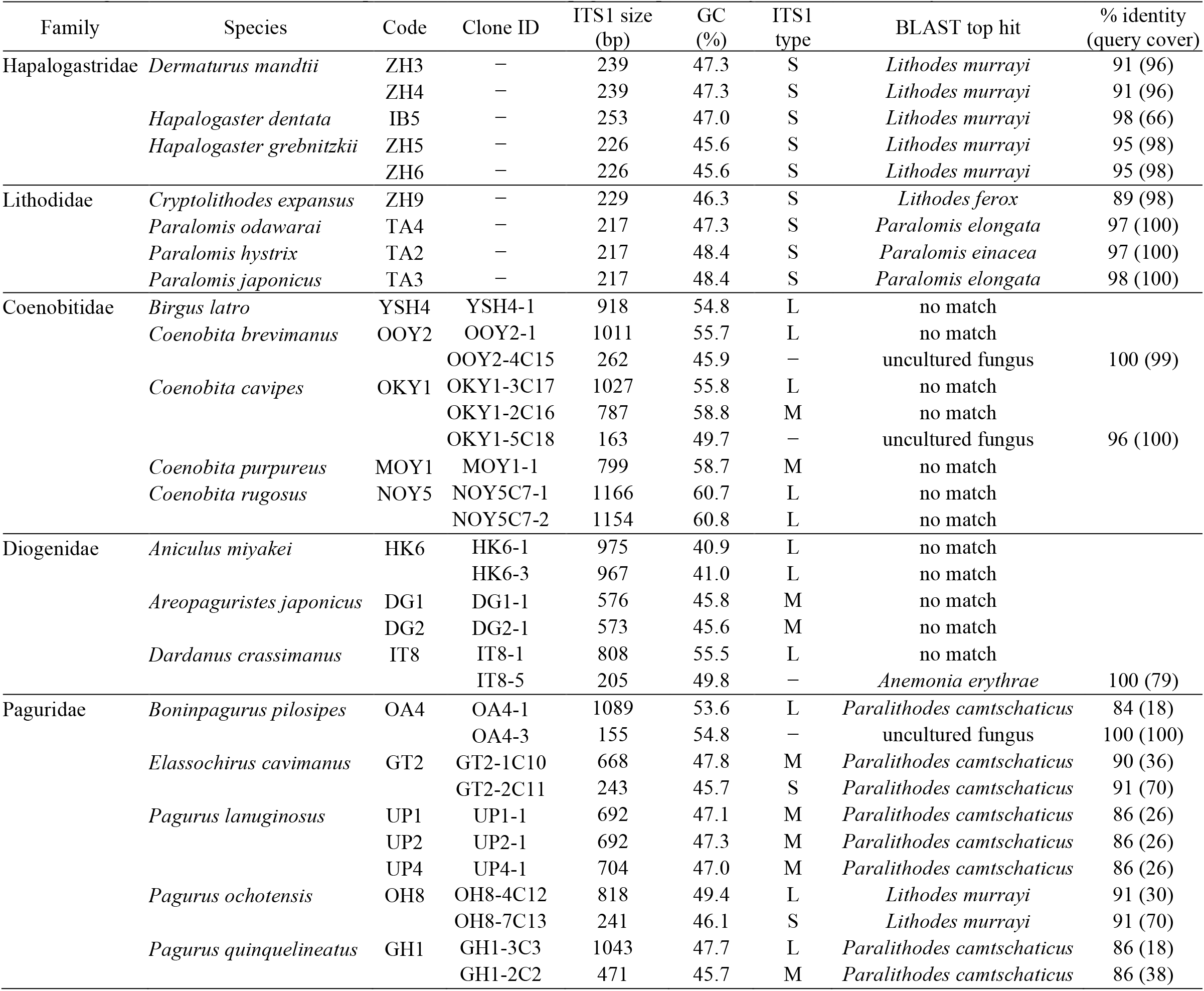
Length and GC content of ITS1 sequences of lithodoid and paguroid species analyzed and the results by BLAST search

| Family | Species | Code | Clone ID | ITS1 size (bp) | GC (%) | ITS1 type | BLAST top hit | % identity (query cover) |
| --- | --- | --- | --- | --- | --- | --- | --- | --- |
| Haplogastriidae | *Dermaturus mandtii* | ZH3 | — | 239 | 47.3 | S | *Lithodes murrayi* | 91 (96) |
|  |  | ZH4 | — | 239 | 47.3 | S | *Lithodes murrayi* | 91 (96) |
|  | *Hapalogaster dentata* | IB5 | — | 253 | 47.0 | S | *Lithodes murrayi* | 98 (66) |
|  | *Hapalogaster grebnitzkii* | ZH5 | — | 226 | 45.6 | S | *Lithodes murrayi* | 95 (98) |
|  |  | ZH6 | — | 226 | 45.6 | S | *Lithodes murrayi* | 95 (98) |
| Lithodidae | *Cryptolithodes expansus* | ZH9 | — | 229 | 46.3 | S | *Lithodes ferox* | 89 (98) |
|  | *Paralomis odawarai* | TA4 | — | 217 | 47.3 | S | *Paralomis elongata* | 97 (100) |
|  | *Paralomis hystrix* | TA2 | — | 217 | 48.4 | S | *Paralomis einacea* | 97 (100) |
|  | *Paralomis japonicus* | TA3 | — | 217 | 48.4 | S | *Paralomis elongata* | 98 (100) |
| Coenobitidae | *Birgus latro* | YSH4 | YSH4-1 | 918 | 54.8 | L | no match |  |
|  | *Coenobita brevimanus* | OOY2 | OOY2-1 | 1011 | 55.7 | L | no match |  |
|  |  |  | OOY2-4C15 | 262 | 45.9 | — | uncultured fungus | 100 (99) |
|  | *Coenobita cavipes* | OKY1 | OKY1-3C17 | 1027 | 55.8 | L | no match |  |
|  |  |  | OKY1-2C16 | 787 | 58.8 | M | no match |  |
|  |  |  | OKY1-5C18 | 163 | 49.7 | — | uncultured fungus | 96 (100) |
|  | *Coenobita purpureus* | MOY1 | MOY1-1 | 799 | 58.7 | M | no match |  |
|  | *Coenobita rugosus* | NOY5 | NOY5C7-1 | 1166 | 60.7 | L | no match |  |
|  |  |  | NOY5C7-2 | 1154 | 60.8 | L | no match |  |
| Diogenidae | *Aniculus miyakei* | HK6 | HK6-1 | 975 | 40.9 | L | no match |  |
|  |  |  | HK6-3 | 967 | 41.0 | L | no match |  |
|  | *Areopaguristes japonicus* | DG1 | DG1-1 | 576 | 45.8 | M | no match |  |
|  |  | DG2 | DG2-1 | 573 | 45.6 | M | no match |  |
|  | *Dardanus crassimanus* | IT8 | IT8-1 | 808 | 55.5 | L | no match |  |
|  |  |  | IT8-5 | 205 | 49.8 | — | *Anemonia erythrae* | 100 (79) |
| Paguridae | *Boninpagurus pilosipes* | OA4 | OA4-1 | 1089 | 53.6 | L | *Paralithodes camtschaticus* | 84 (18) |
|  |  |  | OA4-3 | 155 | 54.8 | — | uncultured fungus | 100 (100) |
|  | *Elassochirus cavimanus* | GT2 | GT2-1C10 | 668 | 47.8 | M | *Paralithodes camtschaticus* | 90 (36) |
|  |  |  | GT2-2C11 | 243 | 45.7 | S | *Paralithodes camtschaticus* | 91 (70) |
|  | *Pagurus lanuginosus* | UP1 | UP1-1 | 692 | 47.1 | M | *Paralithodes camtschaticus* | 86 (26) |
|  |  | UP2 | UP2-1 | 692 | 47.3 | M | *Paralithodes camtschaticus* | 86 (26) |
|  |  | UP4 | UP4-1 | 704 | 47.0 | M | *Paralithodes camtschaticus* | 86 (26) |
|  | *Pagurus ochotensis* | OH8 | OH8-4C12 | 818 | 49.4 | L | *Lithodes murrayi* | 91 (30) |
|  |  |  | OH8-7C13 | 241 | 46.1 | S | *Lithodes murrayi* | 91 (70) |
|  | *Pagurus quinquelineatus* | GH1 | GH1-3C3 | 1043 | 47.7 | L | *Paralithodes camtschaticus* | 86 (18) |
|  |  |  | GH1-2C2 | 471 | 45.7 | M | *Paralithodes camtschaticus* | 86 (38) |

M- and L-type ITS1 sequences (573–1166 bp) were observed in Coenobitidae and Diogenidae, and all types (241–1090 bp) were observed in the family Paguridae. A BLAST search detected no similar sequences for the M- and L-type ITS1 sequences obtained in Coenobitidae and Diogenidae. In contrast, the S-type (241 and 243 bp) and M- and L-types (471 to 1090 bp) in the family Paguridae were highly and partially homologous to the ITS1s of lithodoid species in the database, respectively.

GC content of ITS1 sequences of the family Coenobitidae (54.8 to 60.8%) was significantly higher than the three other categories (Diogenidae, Paguridae, and Lithodoidea) (40.9 to 55.5%) (Mann–Whitney U test, P < 0.005), while no significant difference was observed among those three categories (P > 0.14).

### 3.2 ITS1 sequence variation within species

Similar-sized ITS1 sequences observed within species were highly homologous to each other in the nucleotide sequence. S-type ITS1 sequences (239 bp) determined in two individuals (ZH3 and ZH4) of *Dermaturus mandtii* were identical, as were those (226 bp) determined in two individuals (ZH5 and ZH6) of *Hapalogaster grebnitzkii*. Only one nucleotide substitution was observed between the M-type ITS1 sequences (692 bp) in two individuals (UP1 and UP2) of *P. lanuginosus*.

The alignment of L-type ITS1 sequences (1166 and 1154 bp) detected in two clones (NOY5C7-1 and NOY5C7-2) of *Coenobita rugosus* is shown in Fig. 2. Of the 45 variable sites observed between these sequences, three were nucleotide substitutions and the others were indels associated with a variable number of tandem repeats (VNTR: underlined) responsible for the length difference. Likewise, indels associated with the VNTR were observed to be responsible for the small length difference between the two M-type ITS1 sequences (DG1-1 and DG2-1: 576 and 573 bp) of *Areopaguristes japonicus* (Fig. S1) and among the three M-type ITS1 sequences (UP1-1, UP2-1, and UP4-1: 692 and 704 bp) of *Pagurus lanuginosus* (Fig. S2). In contrast, a length difference not associated with VNTR was observed between the two L-type ITS1 sequences (HK6-1 and HK6-3: 975 and 967 bp) of *Aniculus miyakei* (Fig. S3).

**Fig. 2.**
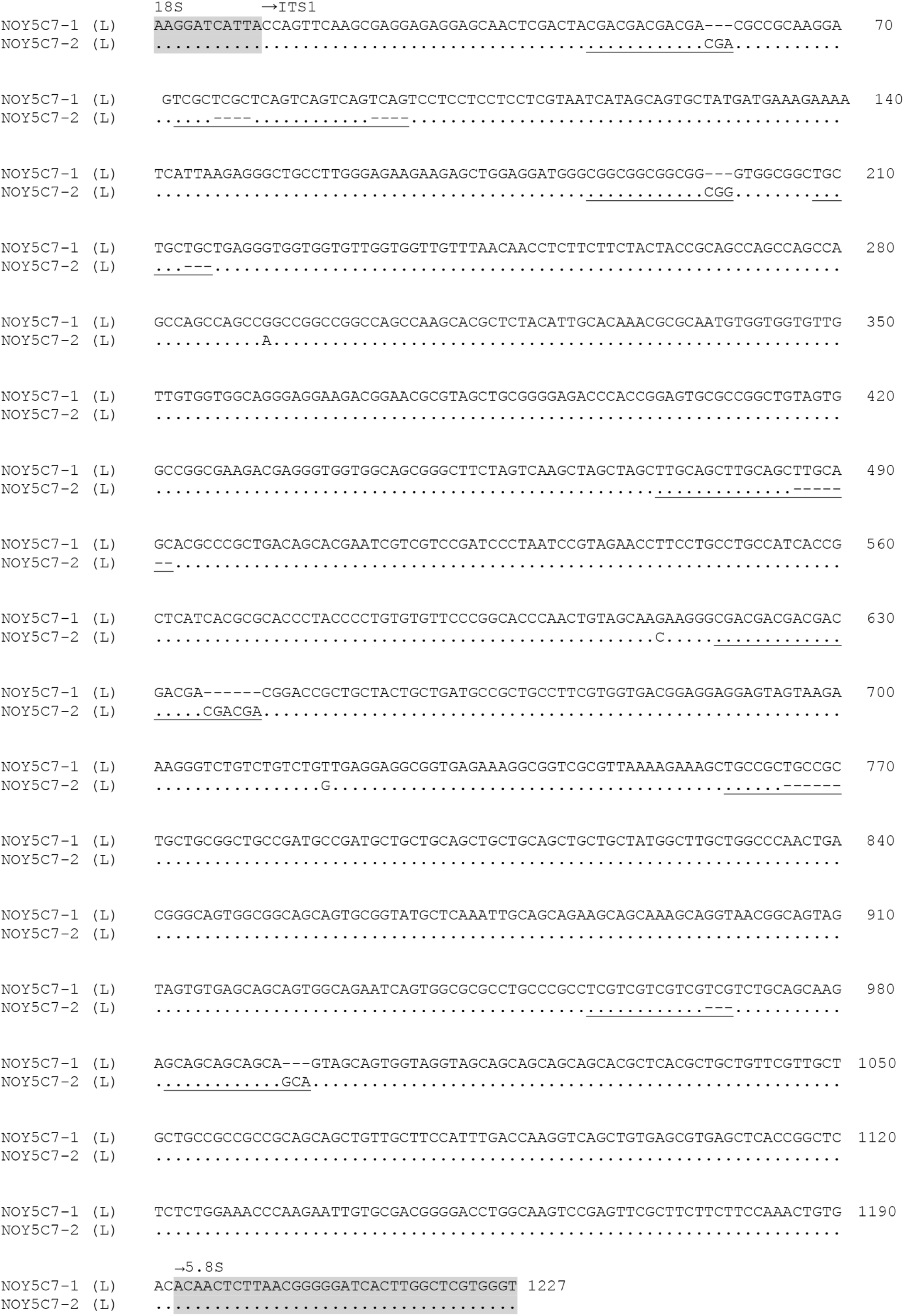
Alignment of two L-type ITS1 sequences (NOY5C7-1 and NOY5C7-2) detected in *Coenobita rugosus*. Dots denote identity to the top sequence, and dashes indicate alignment gaps. Tandem repeats associated with gaps are underlined. Grey shaded regions are 5’ end of 18S rDNA and 3’ end of 5.8S rDNA

Large different-sized ITS1 sequences within species were observed in four species (*Coenobita cavipes, Elassochirus cavimanus, Pagurus ochotensis*, and *Pagurus quinquelineatus*), in which unambiguous sequence alignments between types were obtained in the latter three species. Alignment of the M (GT2-1C10: 668 bp) and S (GT2-2C11: 243 bp) type ITS1 sequences of *E. cavimanus* revealed a large indel (425 bp) (Fig. 3). Such a large indel was also responsible for the significant size difference between L-(OH8-4C12: 818 bp) and S- (OH8-7C13: 241 bp) type ITS1 sequences (577 bp difference) in *P. ochotensis* (Fig. S4) and between L-(GH1-3C3: 1043 bp) and M-(GH1-2C2: 471 bp) type ITS1 sequences (572 bp difference) in *P. quinquelineatus* (Fig. S5). Although similar elements of nucleotide sequences were observed between the L (OKY1-3C17: 1027 bp) and M (OKY1-2C16: 787 bp) ITS1 sequences of *C. cavipes*, alignment was considerably difficult, and no large indels were observed.

**Fig. 3.**
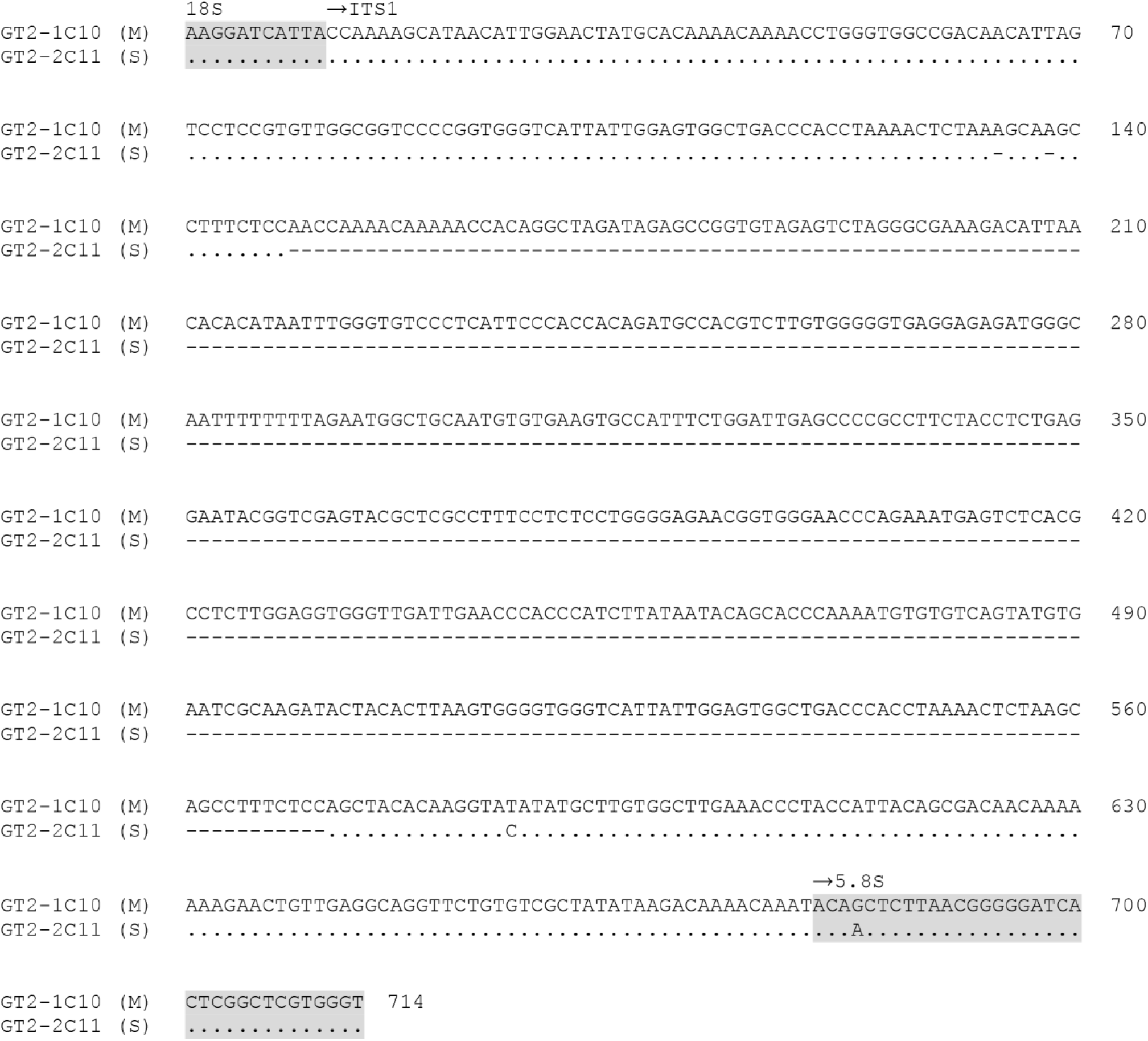
Alignment of M-type (GT2-1C10) and S-type (GT2-2C11) ITS1 sequences detected in *Elassochirus cavimanus*. Type of ITS1 sequence is shown in parenthesis. Dots denote identity to the top sequence, and dashes indicate alignment gaps. Grey shaded regions are 5’ end of 18S rDNA and 3’ end of 5.8S rDNA

### 3.3 ITS1 sequence variation between species and phylogenetic relationships among types and species

As no similar sequences were found in the database for ITS1 sequences of the families Coenobitidae and Diogenidae, these shared no similar sequence elements with the ITS1 sequences of the other families examined in the present study. No appreciable homology was observed between the species of the family Diogenidae.

In the family Coenobitidae, L-type ITS1 sequences of *C. cavipes* (OKY1-3C17: 1027 bp) and *C. brevimanus* (OOY2-1: 1011 bp) were highly homologous, in which VNTRs (underlined) were responsible for the difference in length between them (Fig. 4). Likewise, the M-type ITS1 sequences of *C. cavipes* (OKY1-2C16: 787 bp) and *C. purpureus* (MOY1-1: 799 bp) were highly homologous, in which VNTRs (underlined) were responsible for the length difference between them (Fig. S6). In contrast, the L-type ITS1 sequences of *Birgus latro* (YSH4-1: 918 bp) and *C. rugosus* (NOY5C7-1 and NOY5C7-2: 1166 and 1154 bp) were difficult to align with the L-type ITS1 sequences of *C. brevimanus* and *C. cavipes*. All ITS1 sequences in this family shared conserved regions (underlined) at the 5′ (6 bp) and 3′ region (∼100 bp) (Fig. 5). The phylogenetic tree drawn using the conserved region indicated that the same types of species were more closely related than different types within species (Fig. 6).

**Fig. 4.**
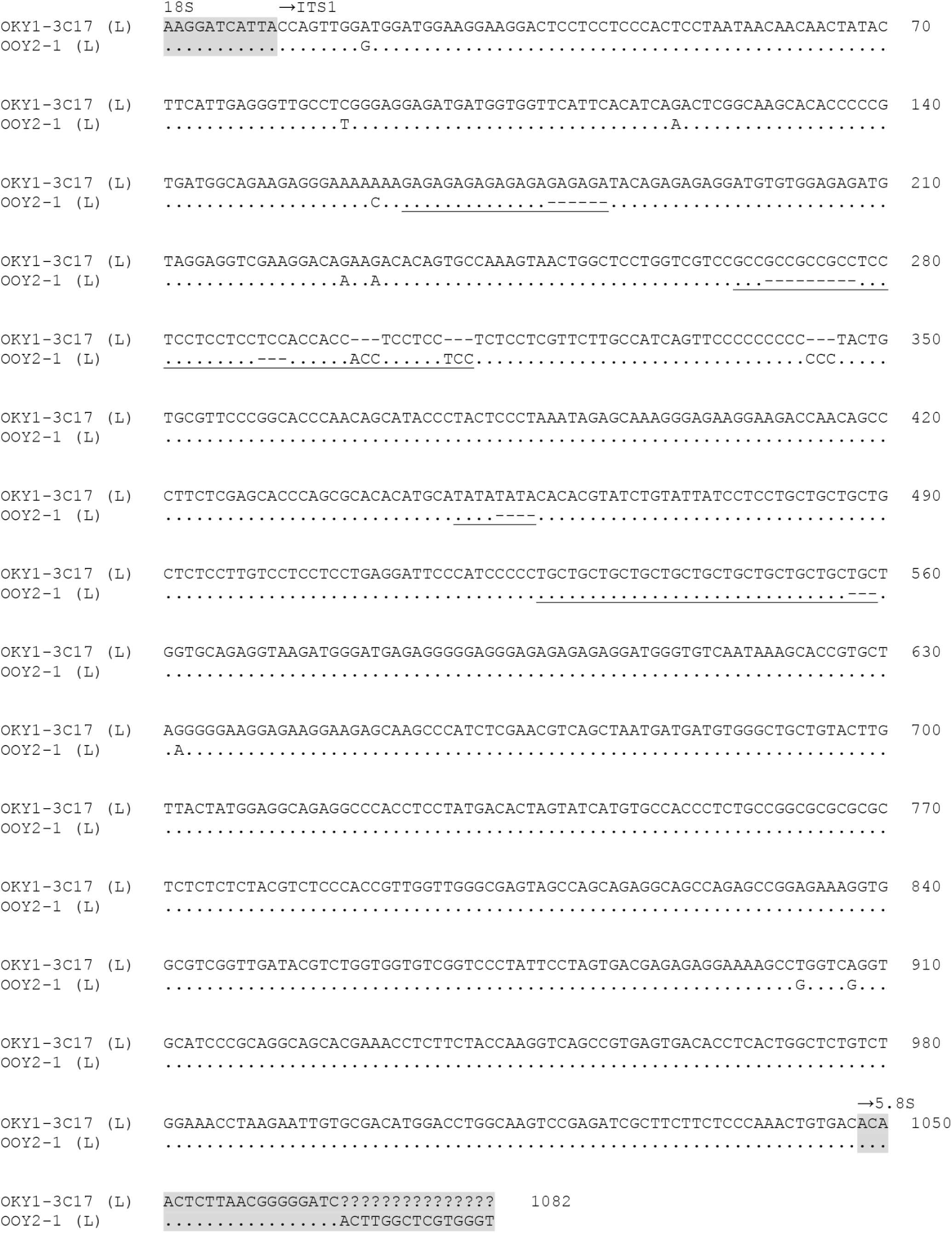
Alignment of L-type ITS1 sequence (OKY1-3C17) of *Coenovita cavipes* and M-type ITS1 sequence (OOY2-1) of *Coenobita brevimanus*. Type of ITS1 sequence is shown in parenthesis. Dots denote identity to the top sequence, and dashes indicate alignment gaps. Tandem repeats associated with gaps are underlined. Grey shaded regions are 5’ end of 18S rDNA and 3’ end of 5.8S rDNA

**Fig. 5.**
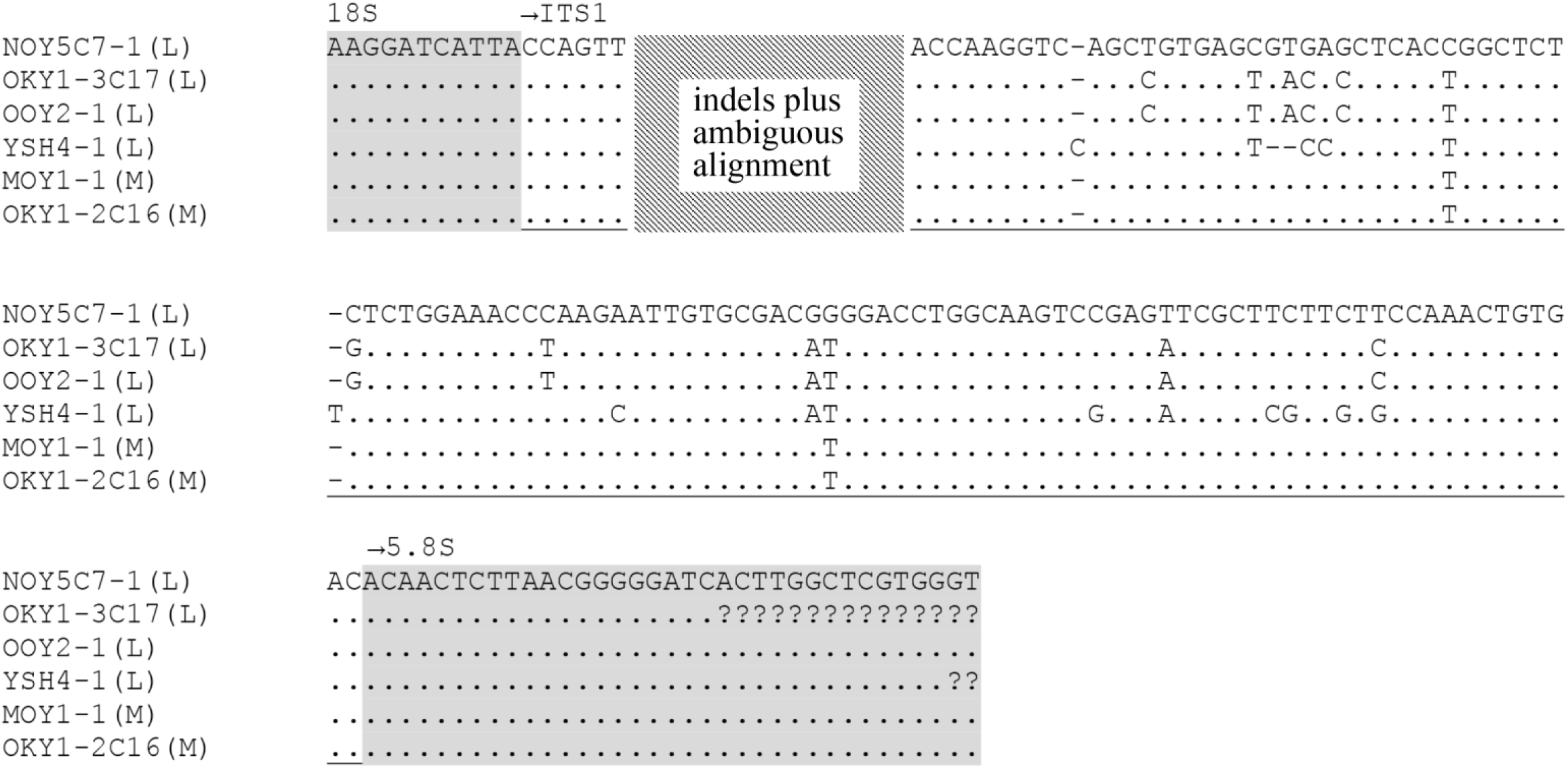
Alignment of relatively conserved c.a. 100 bp 3’ region of four L- and two M-type ITS1 sequences detected in five species of the family Coenobitidae. Type of ITS1 sequence is shown in parenthesis. Dots denote identity to the top sequence, and dashes indicate alignment gaps. Grey shaded regions are 5’ end of 18S rDNA and 3’ end of 5.8S rDNA. See Table 2 for clone ID and species. Relatively conserved region (underlined) was used for phylogenetic analysis

**Fig. 6.**
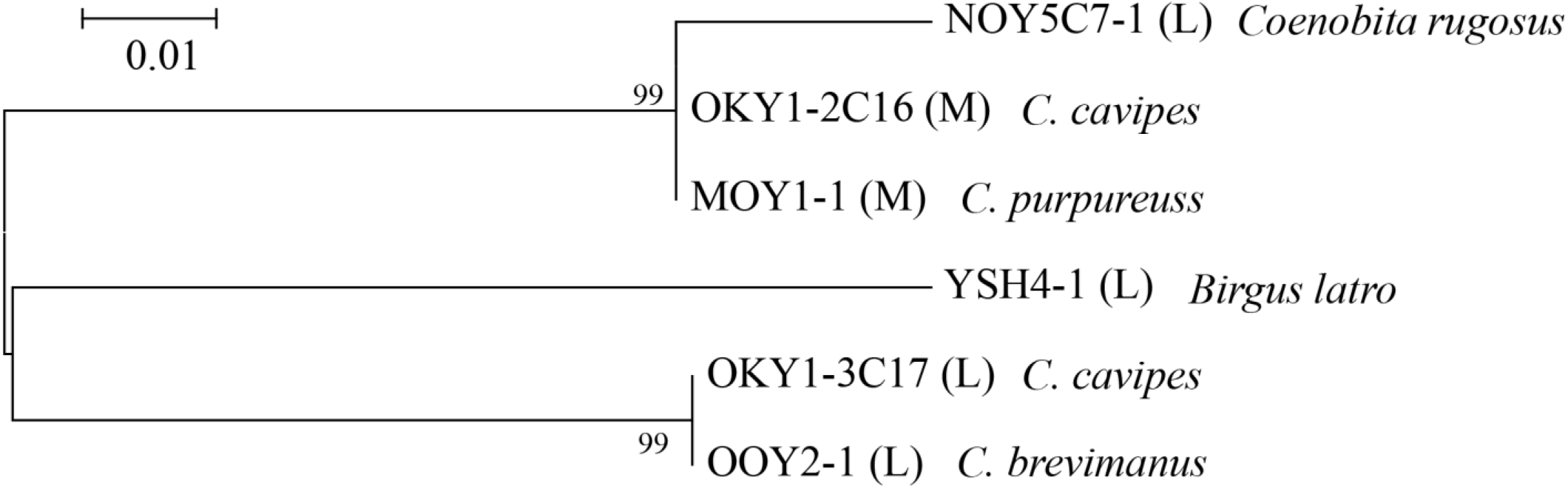
Maximum-likelihood tree (with the best fit JC model) for six sequences of the family Coenobitidae shown in Fig. 8. Type of ITS1 sequence is shown in parenthesis. Bootstrap supports higher than 50% after 1,000 replications are shown at each node

Alignment of ITS1 sequences of the families Hapalogastridae, Lithodidae, and Paguridae is shown in Fig. 7, in which a type-S ITS1 sequence of the king crab (*Paralithodes camtschaticus*), obtained from the database (accession No. AB194389), was incorporated. Although large indels in the central region and multiple VNTRs made the sequence alignment ambiguous, all ITS1 types were observed to share relatively conserved regions (underlined) at the 5′ region (ca. 120 bp) and at the 3′ region (c.a. 120 bp). Furthermore, the M- and L-types shared an c.a. 70 additional bp region (double underline). The phylogenetic tree drawn using the relatively conserved regions among the three types indicated that the different types of the same species were more closely related to each other than to the same types of different species in the family Paguridae (Fig. 8). The same tree topology was obtained when the c.a. 70 additional bp region shared between the M- and L-types was not included. All S-type ITS1s of lithodoid species formed a clade distinct from all types of the family Paguridae; however, the separation between lithodoid families Hapalogarstridae and Lithodidae was not resolved (Fig. 8).

**Fig. 7.**
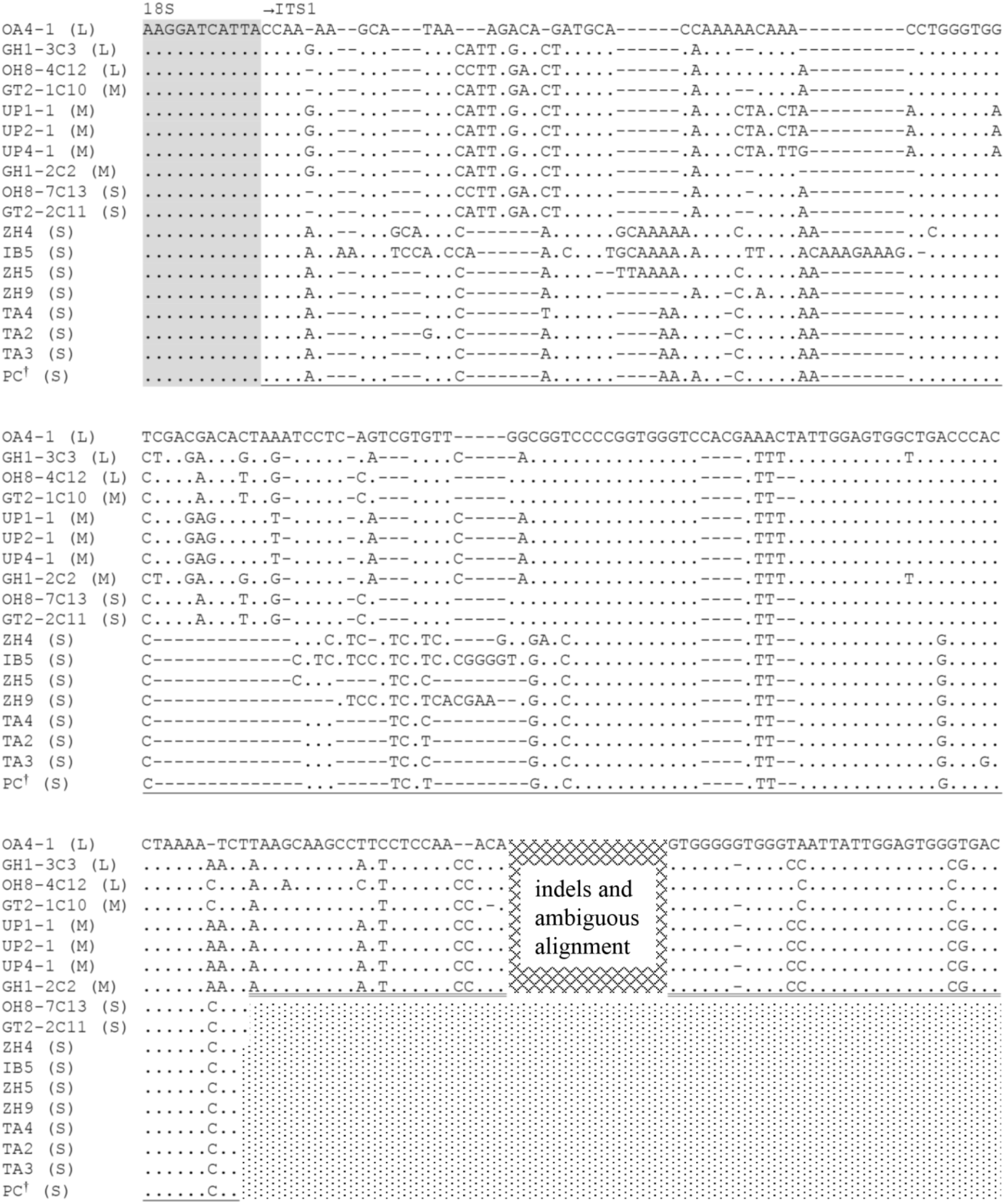
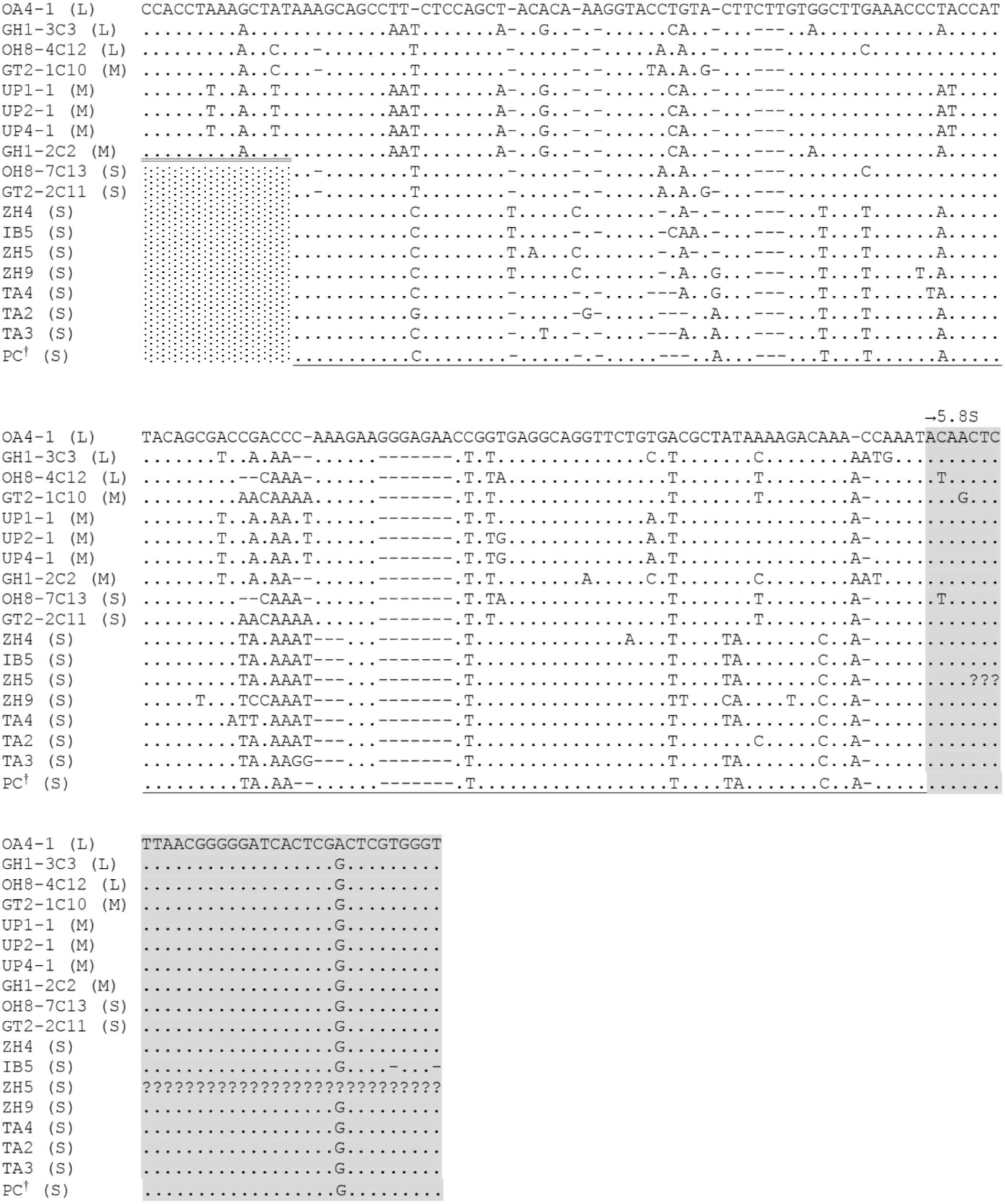
Alignment of three L-, three M- and 10 S-type ITS1 sequences detected in 13 species of the famili palogastridae, Lithodidae and Paguridae. Type of ITS1 sequence is shown in parenthesis. Dots denote entity to the top sequence, and dashes indicate alignment gaps. See Table 2 for clone ID and species. P *ralithodes camtschaticus* ITS1 sequence (AB194389) derived from database. Grey shaded regions are d of 18S rDNA and 3’ end of 5.8S rDNA. Relatively conserved region (underlined) was used for ylogenetic analysis

**Fig. 8.**
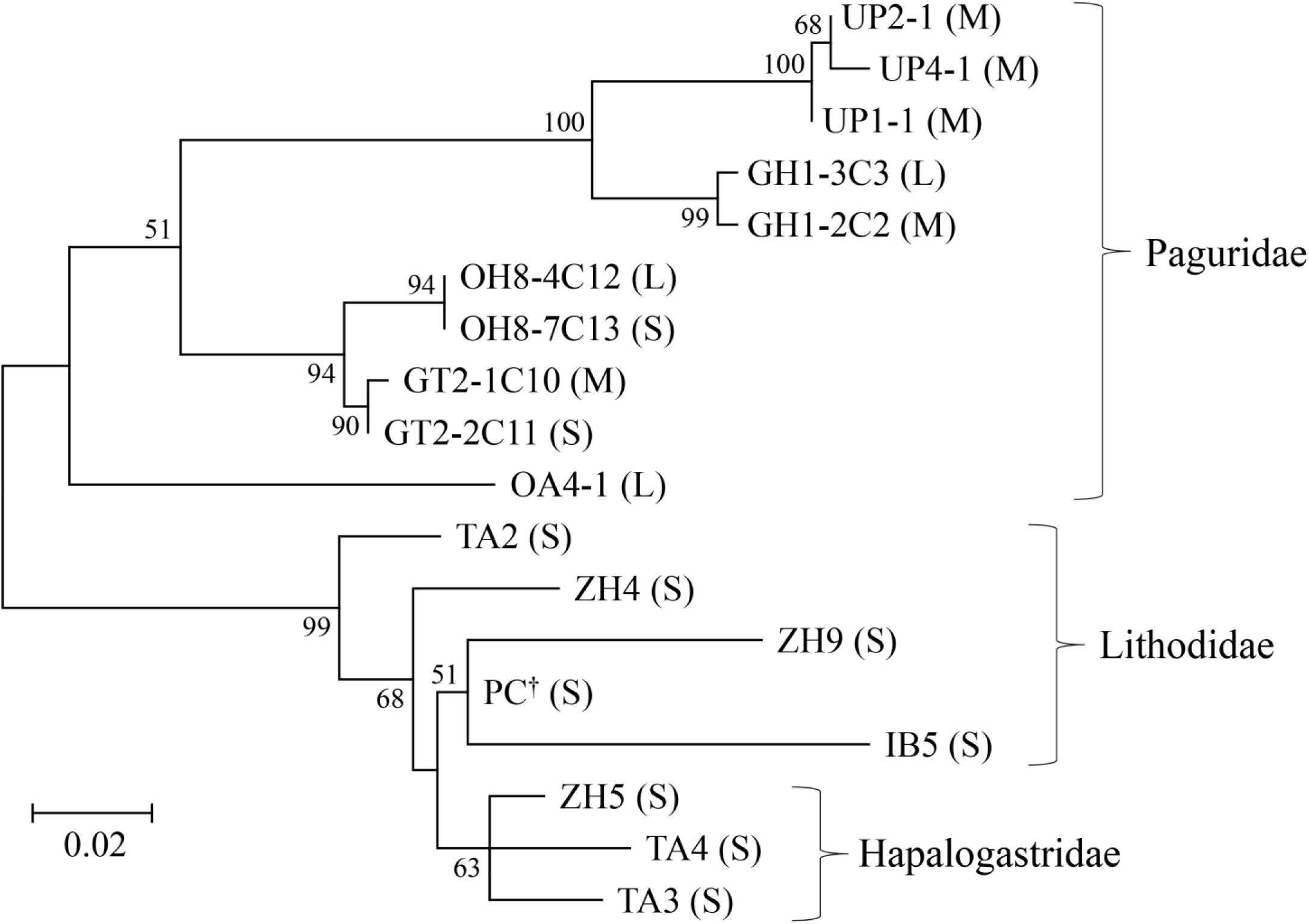
Maximum-likelihood tree (with the best fit JC+G model) for 16 sequences (see Fig. 10) detecte in 13 species of the families Hapalogastridae, Lithodidae and Paguridae. Type of ITS1 sequence is shown in parenthesis. Bootstrap supports higher than 50% after 1,000 replications are shown at each node

## 4 Discussion

### 4.1 Characteristics of ITS1 sequence

The present study is the first to report ITS1 sequences in anomuran species of the paguroid families Coenobitidae, Diogenidae, Paguridae, and the lithodoid family Hapalogastridae. All 54 nucleotide sequences returned by the GenBank database search on the term “Paguroidea internal transcribed spacer 1” as of July 18, 2022, were lithodoid species. The scarcity of ITS1 sequences of paguroid families in the database is probably due to the difficulty of sequence determination by direct nucleotide sequencing, for which amplification of paralogues with different sizes and sequences within individuals may be responsible. Large sequence length differences of ITS1 are sometimes found between closely related (congeneric) species (von der Schulenburg et al. 2001) but are rare within a species (Kauserud and Schumacher 2003). Despite the homogenization force through concerted evolution or molecular drive (Dover 1982; Arnheim 1983), intraspecific or intragenomic variation in ITS1 has been detected in a wide variety of eukaryotes (Harris and Crandall 2000; Ko and Jung 2002; Fairley et al. 2005; Chow et al. 2006; Perez-Barros et al. 2008; Bower et al. 2009; Xu et al. 2009; Hoy and Rodriguez 2013; Gong et al. 2018; Van Wormhoudt et al. 2019). Although VNTRs and large indels are often responsible for intraspecific length variation in ITS1 (Harris and Crandall 2000; Fairley et al. 2005; Chow et al. 2006; Wanna et al. 2006; Van Wormhoudt et al. 2019), the mechanisms underlying these variations may be different. The evolutionary rate of VNTRs may be faster than the pace of concerted evolution, and homogenization by concerted evolution may be much less potent for nuclear ribosomal DNA (nrDNA) on different chromosomes than on the same chromosome (Campbell et al. 1997). Divergent paralogues of ITS1 are frequently detected in anomurans (present study) and crayfish (Harris and Crandall 2000) species, which may be related to their high chromosome counts (Niiyama 1959; Scalici et al. 2010; Mlinarec et al. 2016; Jara-Seguel et al. 2020). However, these paralogues may include multiple functional loci as well as pseudogenes (Buckler et al. 1997). The relatively lower GC content, large intraindividual length, and sequence variation in ITS1 obtained in the present study may suggest some to be pseudogenes of ITS1. At present, however, it is difficult to identify nrDNA pseudogenes, and all criteria for determining pseudogenes are not definitive (Bailey et al. 2003; Gong et al. 2018).

### 4.2 Evolutionary relationships between ITS1 types and between taxa

Our data must be preliminary yet, as partial 18S and 5.8S sequences analyzed were too short to investigate functional or non-functional issues, and the number of clones analyzed per individual was not exhaustive. Assuming that all ITS1 sequences obtained in the present study were functional, we can illustrate an evolutionary relationship among the analyzed anomuran families (Fig. 9). As large nucleotide sequence deletion events are usually more frequent than insertion events (Andersson and Andersson 2001; Lynch 2007), the S-type ITS1 sequences may be descendants of longer ITS1, and the M-type may be from L-type as well. This inference is also supported by the observation that decapod crustacean taxa with short ITS1 (< 400 bp) are rare (Harris and Crandall 2000; Chu et al. 2001; Tang et al. 2003; Wanna et al. 2006; Pérez-Barros et al. 2008; Chow et al. 2009, 2010; Lavery et al. 2014; Van Workhoudt et al. 2019). Our results support the split between the right-handed (Hapalogastridae, Lithodidae, and Paguridae) and left-handed (Coenobitidae and Diogenidae) groups, since the S-type ITS1 was observed only in the former group and the sequences were homologous between species. Coenobitidae may be a relatively recent offshoot among left-handed group. The same ITS1 types between coenobitid species were more closely related than those between different types within species (Fig. 6), indicating that separation between the L- and M-type ITS1s preceded speciation events. Nevertheless, these ITS1 sequences of coenobitid species still shared a c.a. 100 bp relatively conserved sequence in the 3′ region (Fig. 5), whereas no similar sequence element was observed among the species of the other family Diogenidae. This inference requires further investigation, as most of the examined coenobitid species belong to the same genus *Coenobita*. In the right-handed group, all types shared much longer conservative sequences (ca. 240 bp) (Fig. 7) than in the coenobitid species. Only S-type ITS1 was detected in the families Hapalogarstridae and Lithodidae, which formed a clade distinct from all types observed in the family Paguridae (Fig. 8). The S-type ITS1 descendant from longer ITS1s was probably maintained with M- and L-type ITS1s in the right-handed lineage, in which only S-type ITS1 was passed on to the lithodoid lineage, or other types were eliminated in this lineage. However, in the family Paguridae, different types of the same species were more closely related to each other than to the same types of different species (Fig. 8), indicating that separation of these different types occurred after speciation events. This paradox may be solved if large deletions had easily occurred under certain rules, and gain and loss of the S-type ITS1 had repeatedly occurred in each lineage.

**Fig. 9.**
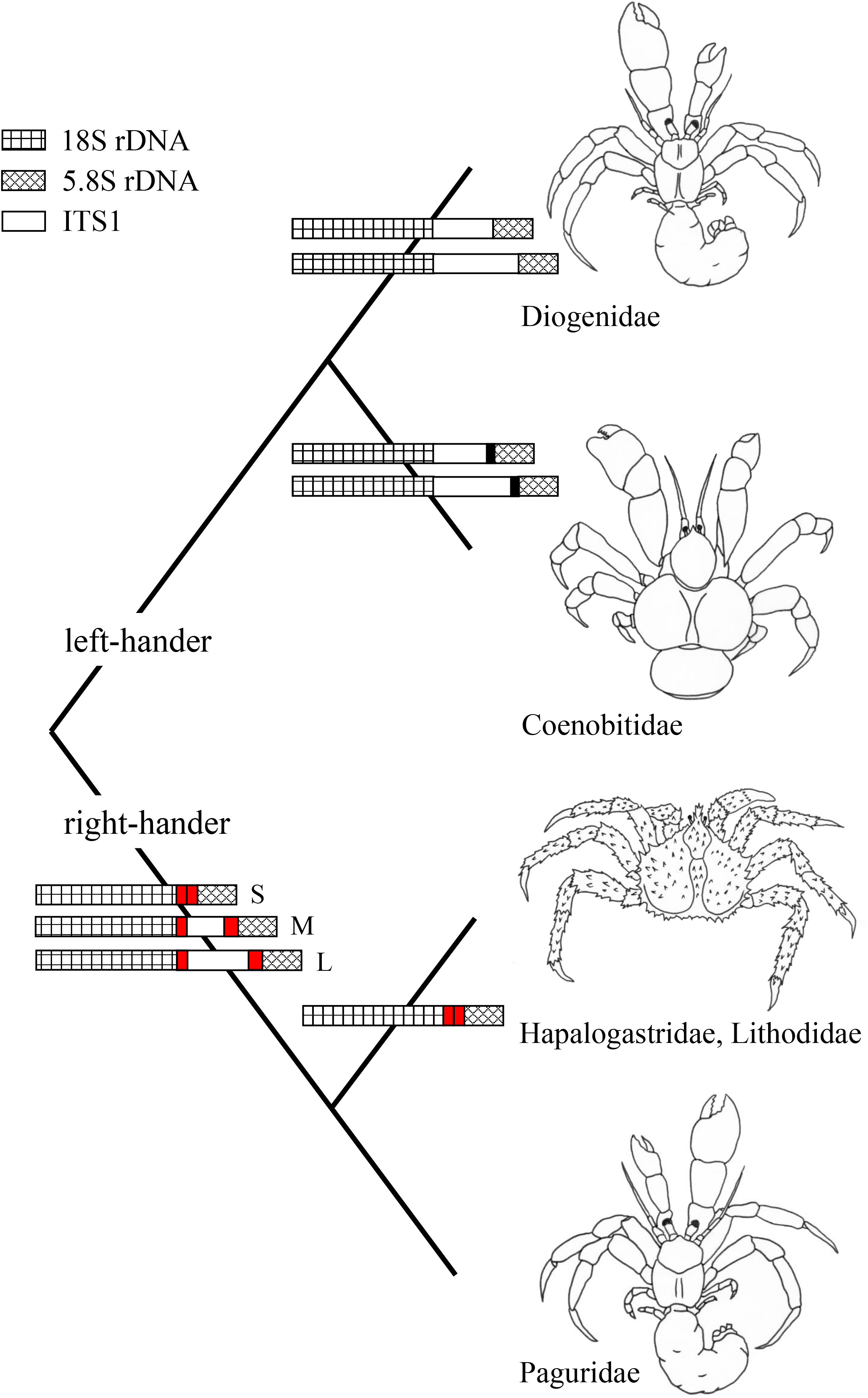
Schematic representation of evolutionary relationship among anomuran families analyzed in the present study based on the size (L: long, M: medium and S: short) and nucleotide sequence variations of ITS1. Black sectors indicate conserved regions of ITS1 sequence among coenobitid species, and red sectors indicate those among litodoid and pagurid species

The evolutionary scenarios mentioned above are concordant with previous molecular phylogenetic studies supporting the “hermit-to-king” hypothesis (Cunningham et al. 1992; Zaklan 2001; Morrison et al. 2002; Ahyong and O’Meally 2004; Tsang et al. 2008, 2011; Ahyong et al. 2009; Bracken-Grissom et al. 2013; Tan et al. 2018, 2019) but contradict the elevation of the lithodoid group to the superfamily rank (McLaughlin et al. 2007) and the nesting of lithodoids within the hermit crab genus *Pagurus* (Cunningham et al. 1992; Ahyong et al. 2009; Schnabel et al. 2011, Bracken-Grissom et al. 2013). Although morphological separation between the families Hapalogastridae and Lithodidae has been proposed (McLaughlin and Lemaitre 1997; McLaughlin et al. 2004, 2007, 2010; Lemaitre and McLaughlin 2009), no clear separation between these families was observed in this study (Fig. 8). Molecular phylogenetic analysis by Bracken-Grissom et al. (2013) and morphological analysis by Zaklan (2001) also indicate that Hapalogastridae is polyphyletic.

This study using multigene family presented supplementary molecular data supporting the “hermit-to-king” hypothesis. Lithodoids are almost certain to harbor little or no L-or M-type ITS1s, since only S-type ITS1 was detected in all seven lithodoid species analyzed in this study, as well as all 19 lithodoid species from the database (as mentioned earlier). In contrast, ITS1 sequence data in the family Paguridae were only available for the five species reported in this study, in which S-type ITS1 of only two species could be analyzed. The heterogeneous distribution of ITS1 types observed among taxa needs further investigation, since ITS1 paralogues of short length and/or lower GC content, including pseudogenes, may be preferentially amplified by PCR (Gong et al. 2016) and all lithodoid ITS1 data were obtained by direct nucleotide sequencing. There may be pagurid species with only S-type ITS1, which would be an important clue for investigating the origin of the king crab.

As mentioned earlier, L- and M-type ITS1s may have been lost in the lithodoid lineage through concerted evolution. An alternative hypothesis is the loss of extensive genomic regions or chromosomes on which L- and M-type ITS1s are clustered. Adaptive phenotypic diversity due to gene loss (Olson 1999; Albalat and Cañestro 2016; Marti-Solans et al. 2021) may have been associated with the morphological transformation of hermit crabs to king crabs.

## 5 Figure legends

**Figure S1**.

Alignment of two L-type ITS1 sequences (HK6-1 and HK6-3) detected in *Aniculus miyakei*. Dots denote identity to the top sequence, and dashes indicate alignment gaps. Tandem repeats associated with gaps are underlined. Grey shaded d regions are 5’ end of 18S rDNA and 3’ end of 5.8S rDNA

**Figure S2**.

Alignment of two M-type ITS1 sequences (DG1-1 and DG2-1) detected in *Areopaguristes japonicus*. Dots denote identity to the top sequence, and dashes indicate alignment gaps. Tandem repeats associated with gaps are underlined. Grey shaded d regions are 5’ end of 18S rDNA and 3’ end of 5.8S rDNA

**Figure S3**.

Alignment of three M-type ITS1 sequences (UP1-1, UP2-1 and UP4-1) detected in *Pagurus lanuginosus*. Dots denote identity to the top sequence, and dashes indicate alignment gaps. Tandem repeats associated with gaps are underlined. Grey shaded d regions are 5’ end of 18S rDNA and 3’ end of 5.8S rDNA

**Figure S4**.

Alignment of L-type (OH8-4C12) and S-type (OH8-7C13) ITS1 sequences detected in *Pagurus ochotensis*. Type of ITS1 sequence is shown in parenthesis. Dots denote identity to the top sequence, and dashes indicate alignment gaps. Grey shaded regions are 5’ end of 18S rDNA and 3’ end of 5.8S rDNA

**Figure S5**.

Alignment of L-type (GH1-3C3) and M-type (GH1-2C2) ITS1 sequences detected in *Pagurus quinquelineatus*. Type of ITS1 sequence is shown in parenthesis. Dots denote identity to the top sequence, and dashes indicate alignment gaps. Grey shaded regions are 5’ end of 18S rDNA and 3’ end of 5.8S rDNA

**Figure S6**.

Alignment of M-type ITS1 sequence (OKY1-2C16) of *Coenovita cavipes* and M-type ITS1 sequence (MOY1-1) of *Coenobita purpureus*. Type of ITS1 sequence is shown in parenthesis. Dots denote identity to the top sequence, and dashes indicate alignment gaps. Tandem repeats associated with gaps are underlined. Grey shaded regions are 5’ end of 18S rDNA and 3’ end of 5.8S rDNA

