## Supplementary Figs. for "Variation of length and sequence of the nuclear ribosomal DNA internal transcribed spacer 1 supports “hermit-to-king” crab hypothesis"

**Journal of Molecular Evolution**

Authors:
Seinen Chow• Katsuyuki Hamasaki • Kooichi Konishi[5]• Takashi Yanagimoto[5]• Ryota Wagatsuma • Haruko Takeyama

Corresponding author:
Seinen Chow
Fisheries Technology Institute


```
           18S        →ITS1
DG1-1 (M)  AAGGATCATTAGAGGTGAAGGAGGATGATGATACACATCTCCCATAAAGAAGGACACTGAACCCTTCGGGTGGGCCTTGG 80
DG2-1 (M)  ................................................................................

DG1-1 (M)  GAGTGAGGTGTTAAGAGTAAAACGAATGTTTTAACTCCCTTTCCTGCTCCCGGCCCCAAGACAAATGAATAATGATACTG 160
DG2-1 (M)  ................................................................................

DG1-1 (M)  CTTCAGCTTTGACAGCTTAGCAGCAAACAACAACAACAGCAGCTCTGGTAATACTTGGTGGGTGAAGAGAGAGGCGGCTC 240
DG2-1 (M)  ...................................---..........................................

DG1-1 (M)  GGTTGATTTACGGATAGTAGTACATGCTACAGAATCTGTTCCCGTTGCATGTTTTGCCTCTCTGAAGGTTTCCTCGCCTC 320
DG2-1 (M)  ................................................................................

DG1-1 (M)  TTACCGTAAGAGTATGCTGCACAATTTGAGGACCACACAGATAACAAATAAGGCGTTGGGGGCAGCCTATGTGTGCAAAA 400
DG2-1 (M)  ................................................................................

DG1-1 (M)  AAGTCCCAATTAATAATAATACATAATAGCACTCCTAGTGGTTGAAGGTCCCACTGCTGGGCCACCACTAAAACTAATAA 480
DG2-1 (M)  ................................................................................

DG1-1 (M)  GACGGTCTATGAGAACTGCTCTTGAAACACCCAATGCCAATTGACGCGAATATGTATGGGACTAACGGGGCGAGTCCCAT 540
DG2-1 (M)  ..................T.............................................................

                                      →ITS1
DG1-1 (M)  TCGCGTAAAAGTATATGAATCTAGAGCACAACTCTTAACGGGGGATCACTCGGCTCGTGGTGT 623
DG2-1 (M)  ....................A..........................................
```

**Fig. S1** Alignment of two M-type ITS1 sequences (DG1-1 and DG2-1) detected in *Areopaguristes japonicus*. Dots denote identity to the top sequence, and dashes indicate alignment gaps. Variable number of tandem repeats associated with gaps are underlined. Grey shaded d regions are 5’ end of 18S rDNA and 3’ end of 5.8S rDNA

```
            18S        →ITS1
UP1-1 (M)  AAGGATCATTACCAAGAAGCATAACATTGGCACTATGCACAAAACTAACTAACTGGGTGACCGAGAGCACTATTCCTCAA 80
UP2-1 (M)  ................................................................................
UP4-1 (M)  ................................................T.G.............................

UP1-1 (M)  TGTTCACGGTCCCCGGTGGGTCATTTTATTGGAGTGGCTGACCCACCTAAAAAACTAAAGCAAGCCATTCTCCAACCACA 160
UP2-1 (M)  ................................................................................
UP4-1 (M)  ................................................................................

UP1-1 (M)  CAAAACTACTACTACTA------------CAACAACAGGCTAGATAGAGCCAGTATAGAGGAGTCTAGGGCAAAAGACCT 240
UP2-1 (M)  .................------------...................................................
UP4-1 (M)  .................CTACTACTACTA...................................................

UP1-1 (M)  CAACACACAATAACCATAATGTTGACCTGTGCCTCATTCCCACTTGGGGGGATGAGAAGGGGAAGGCCAAAATCAGAGTC 320
UP2-1 (M)  ................................................................................
UP4-1 (M)  ................................................................................

UP1-1 (M)  GGTCAAGGCTGTTCTGATTGGGTTCCAATGCGTGAGTGAGAGTCTGGATGAGCCCCGCTTTCTACCTTGAGAGAGGAGAA 400
UP2-1 (M)  ................................................................................
UP4-1 (M)  ................................................................................

UP1-1 (M)  CGGTGGGAAACCAGGCTCGCTGTTATTACAACACGGTAACTAGCACACGCCCCTTGGAGTTGGGTTATTACGGCCCAGCC 480
UP2-1 (M)  ................................................................................
UP4-1 (M)  ................................................................................

UP1-1 (M)  ATCCTAGCTATATGCCGCAGCCAGCATTATGTGTGTGTTTTGTTGTATTGTATTTGTATTGTATTGTGTGGGGTGGGCCA 560
UP2-1 (M)  ................................................................................
UP4-1 (M)  ................................................................................

UP1-1 (M)  TTATTGGAGTGGCGGACCCACCTTAAACTTTAAAGCAGCCAATCTCCAGCAACGCAAGGTACCCATACTTGTGGCTTGAA 640
UP2-1 (M)  ................................................................................
UP4-1 (M)  ................................................................................

                                                                                 →5.8S
UP1-1 (M)  ACCCATCCATTACAGCGTCCAAAACTAAAGAACTGTTGAGGCAGGTTCTGAGTCGCTATAAAAGACAAAACAAATACAAC 720
UP2-1 (M)  ....................................G...........................................
UP4-1 (M)  ....................................G...........................................

UP1-1 (M)  TCTTAACGGGGGATCACTCGGCTCGTGGGT 750
UP2-1 (M)  ..............................
UP4-1 (M)  ..............................
```

**Fig. S2** Alignment of three M-type ITS1 sequences (UP1-1, UP2-1 and UP4-1) detected in *Pagurus lanuginosus*. Dots denote identity to the top sequence, and dashes indicate alignment gaps. Variable number of tandem repeats associated with gaps are underlined. Grey shaded d regions are 5' end of 18S rDNA and 3' end of 5.8S rDNA

```
            18S         →ITS1
HK6-1 (L)  AAGGATCATTACCATTTCATTCGTGTTGTTGGATCTCTTGATATTACCAACGGCAACTTTGTCTGCTTGTTGTTGTTTGA 80
HK6-3 (L)  ................................................................................

HK6-1 (L)  TCCAAAAAAAAAAAGTAAAATAATAGGGTTGCCTTAGGATAATGCACACAAAAACATGCCTGGAAAAGTAAAGAGTACCAT 160
HK6-3 (L)  ..........---...................................................G...........GT..

HK6-1 (L)  TCCAGTTATTGAATAAAGCAAACCAGCGAGTATGGCTGTTTGTCATGTCCAGTACCTACCTATTTGACATCAATAACTAC 240
HK6-3 (L)  ...............................................................................G

HK6-1 (L)  ACTCTCTTTGCTACGGGGGTAAAGTTTTAAATTGTACCCTCACCTCCCTGCTTGGTCGTCCTGTGTGTTCCGGGCACTAA 320
HK6-3 (L)  ..............T........................................................T........

HK6-1 (L)  AGTACAAGGCTTCTGATTTCCGTTTGTTGTTTCTTCTGATTCTTCCCATTCTTGTCTCTTATGAGGGCAAGAAAACTATG 400
HK6-3 (L)  .....................................................C.....A....................

HK6-1 (L)  AAAAAACTAAAAAAGAA-GAAAAAAAAAAAAAAAAAAAGGAAAGAAAGACTGGAAAAAGTGTATGGGTAGAGGTTACAGTCT 480
HK6-3 (L)  ..G..............A...........T------............................................

HK6-1 (L)  GTGGCAAGGTGCTCTCACTGTATCACTCTCTCTCTCTACTTTACTTCCTGGTAAGAAGCCATTTCGTATAGAATAAGTTG 560
HK6-3 (L)  ................................................................................

HK6-1 (L)  TCGTCCCTCTCCCTTCACCTAGAGTAGTAGTCTGTACTACTTTGTTTGTTGTGGTGTTGGTGTCTGGTGGACAGACAGCT 640
HK6-3 (L)  ................................................................................

HK6-1 (L)  AGGAATACTGGAGGTATTACAGCTGGTCATGCTGCCTCTGTCGTTTTTTGCGTCCTCACTCGTCACGAGTCTTTGAGGAT 720
HK6-3 (L)  ................................................................................

800
HK6-1 (L)  TGTGTTGCCAAAGTTGTTTTAATTAACCTTCCCAAAGCTAGGGGAAAAAGAAATAGTGCATGTGGCCAAAGCTGTTTTAG 800
HK6-3 (L)  ................................................................................

HK6-1 (L)  TAGTAGACTACAGCCTTCTGTTCGCCTCCATTTTGAGGTTAGTGGTGGCGGTAGTGGCTTAAAATAGTCTGGTGTCCATG 880
HK6-3 (L)  ..................................................................A.............

HK6-1 (L)  TATTTTTACTTGCCTATTATGTCGGTCTGTTCGATTCTCGAGACCAACTTATTGTGCGACATGGACCTGGCAAGTCTGTT 960
HK6-3 (L)  .........................................A......................................

                                      →ITS1
HK6-1 (L)  TGCTTATTAAGTTTCCAAAACTATGACACAACTCTTAACGGGGGATCACTTGGCTCGTGGGT 1022
HK6-3 (L)  ..............................................................
```

**Fig. S3** Alignment of two L-type ITS1 sequences (HK6-1 and HK6-3) detected in *Aniculus miyakei*. Dots denote identity to the top sequence, and dashes indicate alignment gaps. Simple indels are responsible to length difference. Grey shaded d regions are 5’ end of 18S rDNA and 3’ end of 5.8S rDNA.

```
              18S       →ITS1
OH8-4C12 (L)  AAGGATCATTACCAAAAGCATAACCTTGGAACTATGCACAAAAAACAAAACCTGGGTGGCCGACAACATT 70
OH8-7C13 (S)  ............................................-.........................

OH8-4C12 (L)  AGTCCTCCGTGTTGGCGGTCCCCGGTGGGTCATTATTGGAGTGGCTGACCCACCTAAAACTCTAAAACAA 140
OH8-7C13 (S)  .................................................................G-..-

OH8-4C12 (L)  GCCCTTCTCCAACCACAAACAAAAAACCCCACAGGCTAGATAGAGCCGGTGTAGAGTCTAGGGCGAAAGA 210
OH8-7C13 (S)  ...T......------------------------------------------------------------

OH8-4C12 (L)  CATTAACACACATCATTTGGGTGTCCCTCATTCCCACCACAGATGCCATGTTTTCCGTACCTGGAGTTGG 280
OH8-7C13 (S)  ----------------------------------------------------------------------

OH8-4C12 (L)  GGGGGTTAGACTCGATTCTGTCGGGGCCTCTTATGCGCAGGACGGACTGGGAGCAAAGCGTTTGCAAAAG 350
OH8-7C13 (S)  ----------------------------------------------------------------------

OH8-4C12 (L)  AACATTCGCGATGGGCAGTCTCCCAGACAAAAAACCCTCGACTTACTAAGAGGGGCTGTCTTGTGGGGGT 420
OH8-7C13 (S)  ----------------------------------------------------------------------

OH8-4C12 (L)  GAGGAGAGATGGGCATAAATACAGAATGGCTGCAATGCGTGAGGTACAAGTCTGTGGATGAGCCCCGCCT 490
OH8-7C13 (S)  ----------------------------------------------------------------------

OH8-4C12 (L)  TCAACCTCTGAGGAATACGGTCGAGTACGCTCGCTTTTCCTCTCCTGGGGGGAACGGTGGGAACCCAGAC 560
OH8-7C13 (S)  ----------------------------------------------------------------------

OH8-4C12 (L)  TTGAGTGGCATGCCTCTTGGAGTTGGGTTGATAGAACCCTCCCATCTTATATTACAGCAGCCAAAATGTG 630
OH8-7C13 (S)  ----------------------------------------------------------------------

OH8-4C12 (L)  TGTCAGTATGTGAATTGCAAGATACTACACTTAAGTGGGGGTGGGTCATTATTGGAGTGGCTGACCCACC 700
OH8-7C13 (S)  ----------------------------------------------------------------------

OH8-4C12 (L)  TAAAACTCTAAGCAGCCTTTCTCCAGCTACACAAGGTACATATACTTGTGGCTTCAAACCCTACCATTAC 770
OH8-7C13 (S)  ------------------------..............................................

                                                                         →5.8S
OH8-4C12 (L)  AGCGACCAAAAAAGAACTGTAGAGGCAGGTTCTGTGTCGCTATATAAGACAAAACAAATATAACTCTTAA 840
OH8-7C13 (S)  ......................................................................

OH8-4C12 (L)  CGGGGGATCACTCGGCTCGTGGGT  864
OH8-7C13 (S)  ........................
```

**Fig. S4** Alignment of L-type (OH8-4C12) and S-type (OH8-7C13) ITS1 sequences detected in *Pagurus ochotensis*. Type of ITS1 sequence is shown in parenthesis. Dots denote identity to the top sequence, and dashes indicate alignment gaps. Grey shaded regions are 5’ end of 18S rDNA and 3’ end of 5.8S rDNA

```
                  18S        →ITS1
GH1-3C3 (L)   AAGGATCATTACCAAGAAGCATAACATTGGCACTATGCACAAAACAAACTGGGTGGCTGAGAACAGTAGT   70
GH1-2C2 (M)   ......................................................................

GH1-3C3 (L)   CCTCAATGTTCACGGTCCCCGGTGGGTCATTTTATTGGAGTGGTTGACCCACCTAAAAAACTAAAGCAAG   140
GH1-2C2 (M)   ......................................................................

GH1-3C3 (L)   CCATTCTCCAACCACACAAAACAACAGGTTAGATAGAGCCGGTATCGAAGAGTCTAGGGCAGAAGACCTT   210
GH1-2C2 (M)   ...................---------------------------------------------------

GH1-3C3 (L)   AACACACAAGAACCCTAATGTTGAGCTGTGCTTCATTCCCACTTGTGGGGATGAGAACATTAGGCGTGTG   280
GH1-2C2 (M)   ----------------------------------------------------------------------

GH1-3C3 (L)   TTTGTTGAAACAACAGGCTAGATAGAGCCGGTATAGAACTGCTATCAGACTGATGGTGAGAGAGTTTGGA   350
GH1-2C2 (M)   ----------------------------------------------------------------------

GH1-3C3 (L)   TGAGCCCCGCCTTCTACCTTTCGAGGGGAACGGTAGGAGACCAGACGAAATGCTATCAGACTGATGGACA   420
GH1-2C2 (M)   ----------------------------------------------------------------------

GH1-3C3 (L)   GTCTAGTCTAGGGCGAAAGACTTTAACACACAGGAACCATGATGTTGACCTGTGCCTCATTCCCACTTGT   490
GH1-2C2 (M)   ----------------------------------------------------------------------

GH1-3C3 (L)   GGGGATGAGAAGGGGAAGGCCAAATCTGATAAACGGTTACGGCTGTTTTTGATTGGACTCCAATGCGTGA   560
GH1-2C2 (M)   ----------------------------------------------------------------------

GH1-3C3 (L)   GTGAAAGTCGGGATGAGCCCTGCCTTCTACCTTGAGAGGGGAACGGTGGGAACCAGGCTTGCTGTTATTA   630
GH1-2C2 (M)   ----------------------------------------------------------------------

GH1-3C3 (L)   CAAAACGGTAACACACTCATGCCCCTTGGAGTTGGGTTATTATGACCCAGCCATCCTAGCTGTATGCCGC   700
GH1-2C2 (M)   ----------------------------------------------------------------------

GH1-3C3 (L)   AGCCCAACATTATGTGTGTGTTTGTTGGAACAACAGGCTAGATAGAGCCGGTATAAAACTACTATTGGAC   770
GH1-2C2 (M)   ----------------------------..........................................

GH1-3C3 (L)   TGATGGTGAGAGAGTTTGGATGAGCCCTGCCTTCTACCTTTCGAGGGGAACGGTGGGAGACCAGACGAAA   840
GH1-2C2 (M)   ......................................................................

GH1-3C3 (L)   TGCTATCGGACTGATGAACAGTAGAAAGACCTTGTTGTTATGATGTGTGGGGTGGGCCATTATTGGAGTG   910
GH1-2C2 (M)   ......................................................................

GH1-3C3 (L)   GCGGACCCACCTAAAACTATAAAGCAGCCAATCTCCAGCAACGCAAGGTACCCATACTTGTAGCTTGAAA   980
GH1-2C2 (M)   ......................................................................

GH1-3C3 (L)   CCCAACCATTACAGCGTCCAAAAAAAGAACTGTTGAGGCAGGTTCTGCGTCGCTATACAAGACAAAAATG   1050
GH1-2C2 (M)   .........................................A...........................A

                 →5.8S
GH1-3C3 (L)   AATACAACTCTTAACGGGGGATCACTCGGCTCGTGGGT   1088
GH1-2C2 (M)   ......................................
```

**Fig. S5** Alignment of L-type (GH1-3C3) and M-type (GH1-2C2) ITS1 sequences detected in *Pagurus quinquelineatus*. Type of ITS1 sequence is shown in parenthesis. Dots denote identity to the top sequence, and dashes indicate alignment gaps. Grey shaded regions are 5' end of 18S rDNA and 3' end of 5.8S rDNA

```
               18S        →ITS1
OKY1-2C16 (M)  AAGGATCATTACCAGTTAAGTTAACACGACGACAGTCGTCCTAGTAATACTAAAAATGATGAGGAGGGAT   70
MOY1-1 (M)     ......................................................................

OKY1-2C16 (M)  GCCTTGGGAGAAGAGCAGCAGCAGCAGCAGCAGCTAAGCACGCTCAGCATTGCACAAACAACGCGCAATG  140
MOY1-1 (M)     .........................---------....................................

OKY1-2C16 (M)  TTGTGGCGGCAGGGAGGAAGACGGAACGCATAGCTGCGGGGAGACCCACCGGAGTGCGCCGGCTGTAGTG  210
MOY1-1 (M)     ......................................................................

OKY1-2C16 (M)  GCCGGCGAAGACGAGGGTGGTGGCAGTGGGCAGTGGTAGGAGTAGTAGTAAGCAGCAGCTTGCACGCCCG  280
MOY1-1 (M)     ......................................................................

OKY1-2C16 (M)  CTGACAGCACGAATCGTTGTCCGATCCCTAATCCGTAGAACCTTCCTGCCTGCCGTCACGCGCGGCGCAC  350
MOY1-1 (M)     ......................................................................

OKY1-2C16 (M)  CCTCCTGTGCGTTCCCGGCACCCGACAACGACAGACAGACAGACAGACAGA------------------C  420
MOY1-1 (M)     ...................................................GATAGACAGACGGACAGA.

OKY1-2C16 (M)  CACCGCTTCTATACCGTTGCCTTCATGGTGACGGAGAAGGGGGTGGGGTCGGTCGGTCTACCGAGGTGGT  490
MOY1-1 (M)     ......................................................................

OKY1-2C16 (M)  GAGAATGGCGGTTGCTAATGCTAATGCCGCTGCTGCTGCTGCTGCTGCTGCTGCTGCTGCTATGGATGCT  560
MOY1-1 (M)     ...................................................------------.....C....

OKY1-2C16 (M)  GGCCCAAGTGATGGGCAGTGGCAGCAGCAGTGCGGTATGCTCAAAATGCAGCAGCAGCAGCAGCAGCAGC  630
MOY1-1 (M)     ......................................................................

OKY1-2C16 (M)  AGCAGC---------------GGCTGCCTGCCCACCTTGTCTGTCGTCTCTGGCGGCAGCAGCAGCAGCA  700
MOY1-1 (M)     ......AGCAGCAGCAGCAGC.................................................

OKY1-2C16 (M)  TAGCAGCTGCTTGCTTCCATTTTGACCAAGGTCAGCTGTGAGCGTGAGCTCACTGGCTCTCTCTGGAAAC  770
MOY1-1 (M)     ......................................................................

                                                                            →5.8S
OKY1-2C16 (M)  CCAAGAATTGTGCGACGTGGACCTGGCAAGTCCGAGTTCGCTTCTTCTTCCAAACTGTGACACAACTCTT  840
MOY1-1 (M)     ......................................................................

OKY1-2C16 (M)  AACGGGGGATCACTTGGCTCGTGGGT 866
MOY1-1 (M)     ..........................
```

**Fig. S6** Alignment of M-type ITS1 sequence (OKY1-2C16) of *Coenovita cavipes* and M-type ITS1 sequence (MOY1-1) of *Coenobita purpureus*. Type of ITS1 sequence is shown in parenthesis. Dots denote identity to the top sequence, and dashes indicate alignment gaps. Tandem repeats associated with gaps are underlined. Grey shaded regions are 5’ end of 18S rDNA and 3’ end of 5.8S rDNA
